# Type-I interferons inhibit interleukin-10 signaling and favor type 1 diabetes development in NOD mice

**DOI:** 10.1101/258525

**Authors:** Marcos Iglesias, Anirudh Arun, Maria Chicco, Brandon Lam, Conover Talbot, Vera Ivanova, W. P. A Lee, Gerald Brandacher, Giorgio Raimondi

## Abstract

Destruction of insulin-producing β-cells by autoreactive T lymphocytes leads to the development of type 1 diabetes. Type I interferons (TI-IFN) and interleukin-10 (IL-10) have been connected with the pathophysiology of this disease; however, their interplay in the modulation of diabetogenic T cells remains unknown. We have discovered that TI-IFN cause a selective inhibition of IL-10 signaling in effector and regulatory T cells, altering their responses. This correlates with diabetes development in NOD mice, where the inhibition is also spatially localized to T cells of pancreatic and mesenteric lymph nodes. IL-10 signaling inhibition is reversible and can be restored via blockade of TI-IFN/IFN-R interaction, paralleling with the resulting delay in diabetes onset and reduced severity. Overall, we propose a novel molecular link between TI-IFN and IL-10 signaling that helps better understand the complex dynamics of autoimmune diabetes development and reveals new strategies of intervention.

**Abbreviations:**

ALN: axillary lymph nodes
IL-10: interleukin-10
MFI: mean fluorescence intensity
MLN: mesentheric lymph nodes
NOD: nonobese diabetic mice
PLN: pancreatic lymph nodes
TI-IFN: type-1 Interferons
Tmem: memory T cells
Treg: regulatory T cells

## 1. Introduction

Type 1 Diabetes is a complex autoimmune disease characterized by the progressive destruction of the insulin-producing β-cells in the pancreas by autoreactive T lymphocytes [1]. It is caused by a combination of genetic predisposition and environmental factors. In the past decade, viral infections and the composition of the gut microbiota have gained increasing attention as environmental factors that contribute to the initiation of the disease[2–4] but the mechanisms by which these factors contribute to the activity of diabetogenic T cells remains unknown. It is clear that CD4 T cells are of chief importance in this disease. Nonobese diabetic (NOD) mice, a widely used model of human T1D that spontaneously develop diabetes, are protected from the disease onset when deficient in CD4 T cells [5, 6] and enriched CD4^+^ cells from diabetic donors are able to transfer the disease when administered into NOD-scid/scid recipients [7]. However, the connection between environment and the activity of diabetogenic T cells remains elusive.

Multiple clinical and experimental observations point toward type-I interferons (TI-IFN), essential cytokines for the clearance of viruses, as the mediators that drive a pre-diabetic or susceptible individual toward type 1 diabetes [8]. Relevant examples include: high levels of IFNα detected in the pancreas of diabetic patients [9], absence of autoantibodies able to neutralize IFN-α in the subset of AIRE-deficient (APS1) patients that developed diabetes [10], induction of diabetes in non-autoimmune prone C57BL/6 mice by overexpression of IFN-α in β-cells [11], accumulation of high levels of TI-IFN in NOD mice [12], and delay of disease onset (and decreased incidence) with early blockade of TI-IFN receptor signaling [13]. More recently, Ferreira and colleagues reported that an IFN signature in PBMC of genetically predisposed children was detectable before the appearance of islet-specific autoantibodies [14]. Despite these observations, the mechanism(s) through which TI-IFN promotes T1D remains poorly understood.

The cytokine IL-10 has an essential role in the development of autoimmune pathologies.[15] Previous studies suggested that the low expression of this cytokine in the pancreas mediates the occurrence of diabetes [16] and decreased IL-10 levels in serum of newly diagnosed children with type 1 diabetes has been observed [17]. Monocytes/macrophages have been historically investigated as the main target of this cytokine [18]; however, IL-10 acts also directly on T cells. This has been shown in the context of naïve T cells activation and differentiation [19, 20], in the regulation of effector and memory T cells [21–23] and in the preservation of regulatory T cell function [24, 25].

Here we report a novel effect of TI-IFN that causes a selective inhibition of IL-10 signaling in T cells thereby reducing their capacity to be regulated. This loss of signaling correlates with the development of the disease in NOD mice. This effect is sustained but compartmentalized, manifesting only in T cells of pancreatic (PLN) and mesenteric (MLN) lymph nodes of NOD mice, suggesting a link with the response to the gastric environment in these animals. Importantly, IL-10 signaling in T cells could be partially restored via blockade of TI-IFN signaling, supporting earlier observations on the beneficial effects of transient TI-IFN blockade in NOD mice [13]. Overall, our results reveal a new molecular mechanism involved in the causative process of type 1 diabetes and suggest novel targets for prevention and treatment of T1D.

## 2. Results

### 2.1 Localized defective IL-10 signaling in memory and regulatory CD4 T cells in TI-IFN enriched tissues.

Li et al. reported unexpectedly high levels of IFNα production in the PLN of NOD mice starting in the 2^nd^ week of life that correlated with the presence of CD4 T cells with a transcriptional signature abundant in INF-induced genes [13]. Based on the suspected involvement of IL-10 in disease development, we tested if these TI-IFN-exposed T cells would show any alteration in their response to IL-10. To evaluate signal integrity, we quantified the accumulation of the phosphorylated (active) form of the transcription factor STAT3 (P-STAT3, a key molecule in the IL-10 signaling pathway) in response to *ex vivo* stimulation with IL-10. The response to the pro-inflammatory cytokine IL-6 (that also induces phosphorylation of STAT3) was measured to distinguish between cytokine-specific Vs nonspecific effects of TI-IFN exposure. We compared multiple CD4 T cell subsets: naïve (CD4^+^CD44^low^Foxp3^-^), memory (CD4^+^Foxp3^-^CD44^hi^) and regulatory (CD4^+^Foxp3^+^) T cells residing in PLN, MLN, axillary lymph nodes (ALN), and spleen (each separately), in 4-week-old NOD mice (the reported time of highest accumulation of IFNα; [13]). Independent repeats of these measurements indicated a statistically significant reduction in IL-10 signaling in both Tmem and Treg from PLN and MLN compared to the response of the same T cell subsets in the spleen (Fig. 1A&B) and ALN (not shown). In 4-week-old non-diabetes prone B6 mice, which do not accumulate TI-IFN in pancreatic and mesenteric lymph nodes, Tmem and Treg preserved their ability to fully respond to IL-10 in all lymphoid tissues (Fig. 1A&C). Importantly, this decrement in STAT3 phosphorylation was specific to IL-10 signaling, as the response to IL-6 was unaltered in the T cells of NOD (and B6) mice from all the lymphoid tissues tested (Fig. 1C). Together, these results suggested that in NOD mice there is a selective reprogramming of the signaling for IL-10, actuated specifically in lymphoid tissues shown to accumulate TI-IFN [13].

**Figure 1.**
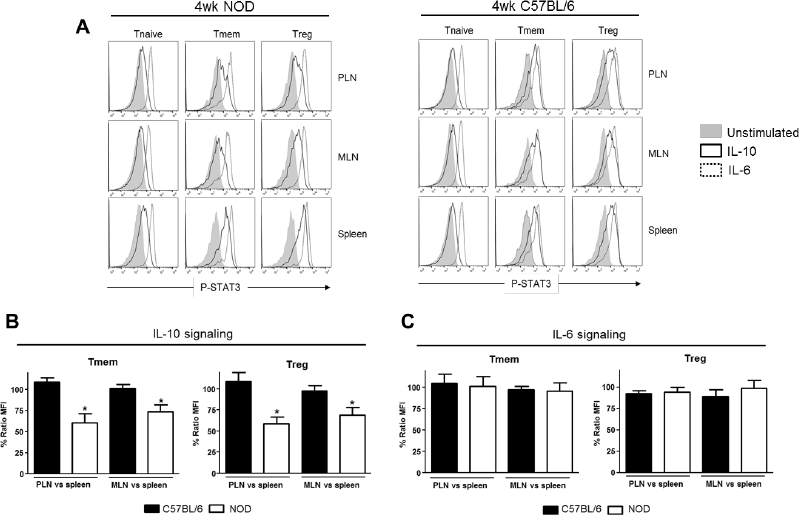
Defective IL-10 signaling in Tmem and Treg cells in pancreatic and mesenteric lymph nodes of NOD mice. Cells of pancreatic lymph nodes (PLN), mesenteric lymph nodes (MLN), and spleens of 4 week old NOD or C57BL/6 mice were either left untreated or stimulated with IL-10 (40 ng/ml) or IL-6 (40 ng/ml) for 20 min. Phosphorylated-STAT3 (P-STAT3) levels induced in CD4 T cell subpopulations (Tnaive: CD4^+^CD44^low^Foxp3^-^, Tmem: CD4^+^CD44^hi^Foxp3-, Treg: CD4^+^Foxp3^+^) were measured by Phospho-flow. (**A**) Representative histograms of PSTAT3 levels in the indicated CD4 T cell subpopulations after IL-10 and IL-6 stimulation. (**B-C**) Cumulative representation of the percentage of P-STAT3 signal induction in PLN/MLN Tmem and Treg after IL-10 (B) or IL-6 (C) stimulation compared to level induced in the splenic populations of the indicated mouse strain. This ratio of MFIs was calculated by comparing the coefficient index of P-STAT3 between two different tissues, considering the levels in spleen as 100% of expression. Data shown in (B) and (C) is the average from n=4 mice per strain and is expressed as % of Ratio MFI±SEM, *p<0.05, paired Student‘s t test.

### 2.2 The impact of TI-IFN on IL-10 signaling is not a genetic characteristic of NOD T cells.

We then tested if this effect was unique to T cells of NOD background, or if bystander exposure of any T cells to unusual levels of TI-IFN could affect their ability to be controlled by IL-10.Bulk T cells from wt B6 mice were exposed to IFN-β (or IFN-α) for 48 hours, and then the levels of P-STAT3 induced by stimulation with IL-10 or IL-6 were quantified via Phospho-flow. Exposure to IFN-β (and similarly to IFN-α, not shown) induced a statistically significant reduction of STAT3 phosphorylation after IL-10 stimulation in Tmem and Treg cells when compared to the response in fresh or mock-treated cells (cultured without IFN-β, to exclude any impact from the culturing conditions) (Fig. 2A&B). Reduction of IL-10 signaling responses were dose-dependent upon IFN-β levels, reaching maximum inhibition at 5 ng/ml of IFN-β (Supplementary Fig. 1). As observed in T cells from NOD mice, the levels of P-STAT3 in response to IL-6 stimulation remained unaltered under all conditions (Fig. 2C), confirming that this effect was not a generalized saturation of the Jak/STAT signaling pathway.

**Figure 2.**
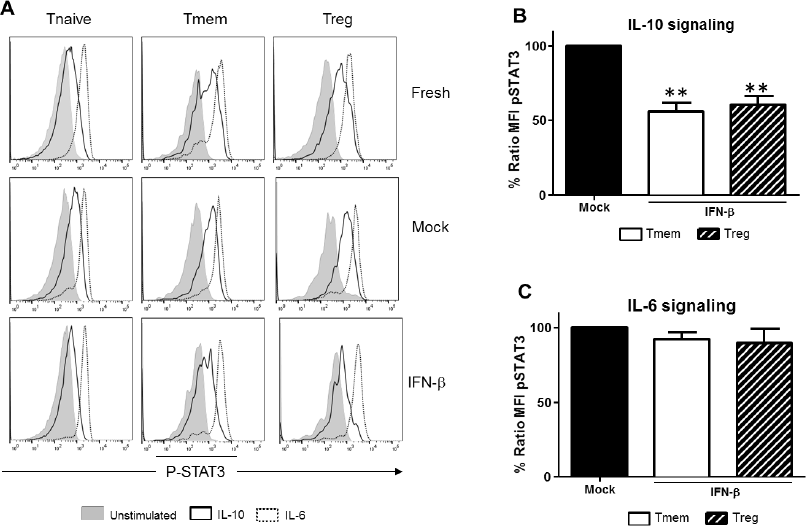
Exposure to IFN-β induces a selective inhibition of IL-10 signaling in CD4 Tmem and Treg irrespective of the strain of origin. Purified T cells from C57BL/6 mice were freshly stimulated or cultured for 48h in complete media with or without IFN-β (5ng/ml), and rested in cytokine-free media for six additional hours. Cells were then either left untreated or stimulated with IL-10 (40 ng/ml) or IL-6 (40 ng/ml) and the levels of P-STAT3 in CD4 T cell subpopulations were measured by phospho-flow as in Figure 1. (**A**) Representative histograms of the MFI levels of P-STAT3 in the different CD4 T cell subpopulations of fresh, mock and IFN-β exposed T cells after IL-10 and IL-6 stimulation. (**B-C**) Graph bars that compare the percentage of P-STAT3 MFI ratio between IFN-β exposed and not exposed (mock) cultured Tmem and Treg cells after IL-10 (B) or IL-6 (C) stimulation. Ratio MFI was calculated comparing the coefficient index of PSTAT3 after stimulation between the two different culture conditions, considering levels in mock cells as 100% of expression. Data from n=6 individual experiments are shown and expressed as % of Ratio MFI±SEM, **p<0.01, paired Student‘s t test.

### 2.3 IFN-β–mediated inhibition of IL-10 signaling alters induction of IL-10 responsive genes

A deeper evaluation of the functional impact that inhibition of IL-10 signaling by TI-IFN has on the modulation of T cells can be done via assessment of its transcriptional impact. However, the transcriptional impact of IL-10 on T cells is unknown. We therefore harnessed the vast knowledge about IL-10 signaling in antigen presenting cells. Taking advantage of data from the most recent publicly available RNAseq analysis of mouse macrophages exposed to IL-10 [26], we selected a pool of 29 genes highly upregulated (> 6σ) (Supplementary Fig. 2A) as initial lead for genes that could be also induced in T cells. Ten of these upregulated genes had known functions in T cells (Supplementary Fig. 2B). We then tested the expression of these genes as a screening panel for further investigation of Treg and Tmem cells responses to IL-10 with or without TI-IFN pre-exposure. To obtain a sufficient and homogeneous number of Tmem, we implemented the previously published “parking method” (Methods) where in vitro-activated T cells are “parked” in congenic Rag^−/−^ mice to generate Tmem [27]. Treg cells were freshly-isolated from unmanipulated animals. Gene transcription analysis showed four genes –*LIGHT, Sphk1, Tarm1* and *2B4* – to be significantly upregulated in Tmem by in vitro treatment with IL-10 (Fig. 3A), and two genes, *Sphk1* and *2B4,* were upregulated in Treg (Fig. 3B). We then analyzed if the induction of these genes was affected by pre-exposure of these cells in vitro to IFN-β. In Tmem, the increased expression of *LIGHT, Sphk1, Tarm1* and *2B4* was completely abrogated (Fig. 3A). mRNA levels of *Sphk1, LIGHT* and *Tarm1* also showed an important decrease in IFN-β exposed Treg, while the expression of *2B4* was not affected (Fig. 3B). These results indicate that the selective inhibition of STAT3 phosphorylation induced by INF-β pre-exposure alters significantly the impact of IL-10-mediated transcription in Tmem and Treg cells.

**Figure 3.**
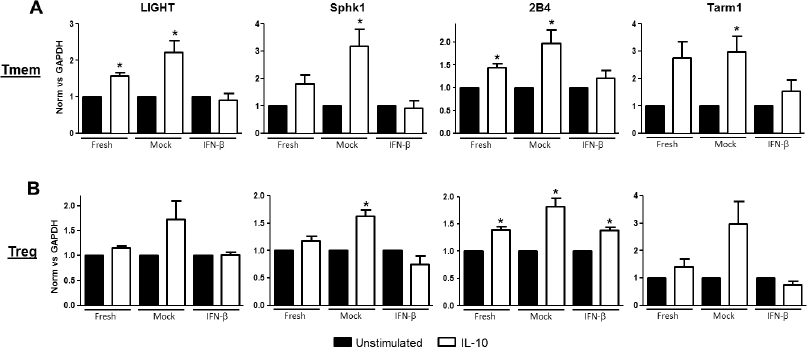
Expression of IL-10 regulated genes in T cells is abrogated after exposure to IFN-β. (A) Freshly purified CD4 Tmem cells (obtained using the “parking method“; see material and methods) and (B) CD4^+^CD25^+^ Treg cells from C57BL/6 mice were left untreated or cultured 48h without (Mock) or with IFN-β (5ng/ml). After an additional 4 hours of resting culture (without stimuli), cells were either left untreated or stimulated for 4 hours with IL-10 (40 ng/ml). Cell were then lysed and mRNA levels of *LIGHT, Sphk1, 2B4* and *Tarm1* genes were measured by qPCR.ΔΔCt method was used to calculate their relative expression and normalized to GAPDH. The graph bars show the fold change between unstimulated and IL-10 stimulated cells in every condition ±SEM where *p<0.01 in paired Student‘s t test was considered with statistically significance. Data shows the average of n=3 independent experiments.

### 2.4 Inhibition of IL-10 signaling requires prolonged exposure to IFN-β, but it is reversible

To explore what length of exposure to IFN-β is required to impact IL-10 signaling in T cells, wt B6 bulk T cells were exposed to IFN-β for different lengths of time and the response to IL-10 and IL-6 in different subsets assessed by Phospho-Flow. In Tmem cells, a 24-hour exposure significantly reduced the levels of IL-10-induced P-STAT3, but a 48-hour exposure was necessary to achieve maximal inhibition (Fig. 4A). In Treg cells, a 24-hour exposure was sufficient to achieve the maximal inhibition of IL-10-induced P-STAT3 signaling (Fig. 4A).We also tested the reversibility of this inhibition. To address this question, after 48 hours of exposure to IFN-β, T cells were washed, rested in cytokine-free media for 24 or 48 hours and the P-STAT3 response to IL-10 or IL-6 was then measured. Within 24 hours of removing IFN-β, Treg recovered their normal P-STAT3 response to IL-10 (Fig. 4B). The recovery of Tmem was slower, showing only partial restoration of the IL-10 signaling even at 48 hours after removing the IFNβ (Fig. 4B). These results indicate that the bystander effect of IFN-β requires a prolonged exposure to instigate inhibition of IL-10 signaling and, with some kinetic differences between Treg and Tmem, a normal P-STAT3 response to IL-10 in T cells can be restored following removal of IFN-β.

**Figure 4.**
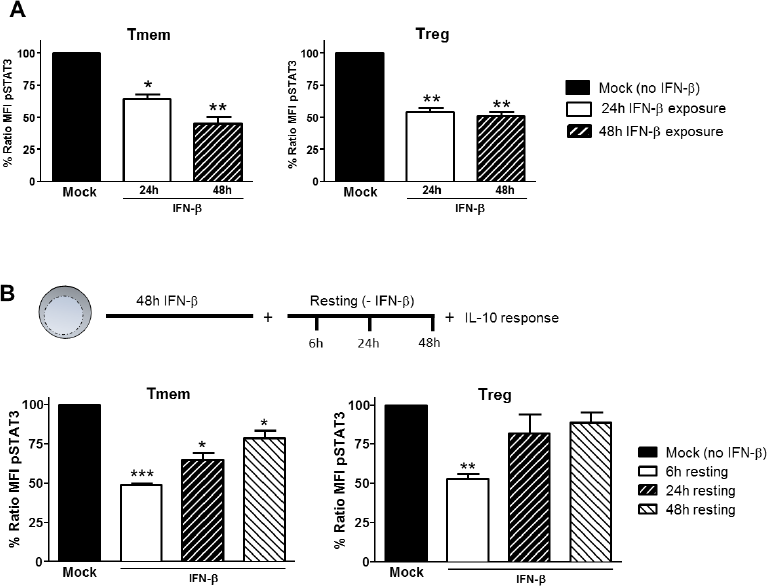
Induction of IL-10 signaling inhibition in T cells requires prolonged exposure to IFN-β but is reversible. (**A**) Purified T cells from C57BL/6 mice were cultured for 24 or 48h with or without IFN-β (5ng/ml), and rested in cytokine-free media for six additional hours. Cells were then left untreated or stimulated with IL-10 (40 ng/ml) and the inhibition of P-STAT3 induction by IL-10 assessed as indicated in Figure 1. (**B**) Purified T cells from C57BL/6 mice were cultured for 48h with or without IFN-β (5ng/ml) and then rested in cytokine-free media for the indicated time (6, 24, 48h). The levels of P-STAT3 in response to IL-10 (40 ng/ml) were then assessed as previously indicated. The ratio MFI (panels A-B) was calculated comparing the coefficient index of P-STAT3 after IL-10 stimulation between IFN-β exposed and mock, considering levels in mock cells as 100% of expression. Data of n=3 individual experiments are shown and expressed as % of Ratio MFI±SEM, *p<0.05, **p≤0.01, ***p≤0.01, paired Student‘s t test.

### 2.5 IFN-β signals through the Jak/STAT pathway to inhibit IL-10 signaling in T cells

TI-IFNs signal through multiple pathways [28], with the activation of the Jak/STAT route considered the most relevant to their antiviral effects. To identify the signaling pathway responsible for the actuation of IFN-β induced perturbations of IL-10 signaling, cells were pre-treated with two small molecule Jak-specific inhibitors, Tofacitinib and Ruxolitinib, prior to and during IFN-β stimulation. Tofacitinib inhibits Jak3 and Jak1, while Ruxolitinib blocks Jak2 and Jak1. After preconditioning bulk T cells with Tofacitinib (25 µM) or Ruxolitinib (5 µM) for 2 hours, IFN-β was added to the cultures for 48 hours. Following incubation and washing, an additional 6-8 hours resting phase allowed the cells to recover their signaling after removal of the Jak inhibitor (Supplementary Fig. 3). The impact on IL-10 or IL-6 signaling was then quantified via phospho-flow. In Tmem, Tofacitinib treatment resulted in a statistically significant preservation of IL-10 signaling and Ruxolitinib had an even stronger effect (Fig. 5A). In Treg, both inhibitors restored IL-10 signaling to the same extent (Fig. 5A), though the cells had to be rested for 8 hours (instead of 6 hours as in the case of Tmem), as their recovery of cytokines signaling after Jak inhibition was slower than in Tmem. Collectively, these results suggest that Jak1, and possibly Jak2, are essential mediators for IFN-β—mediated alterations of IL-10-induced P-STAT3 signaling in T cells.

**Figure 5.**
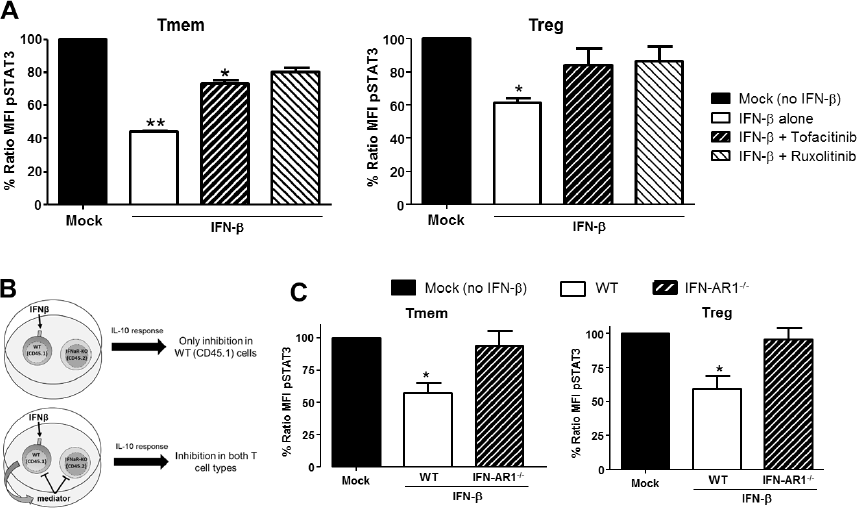
IFN-β signals through JAK-STAT pathway to directly inhibit IL-10 signaling in T cells. (**A**) Impact of JAK-STAT inhibition on the ability of IFN-β to modulate IL-10 signaling. T cells from C57BL/6 mice were exposed to Tofacitinib (25µM) or Ruxolitinib (5µM) for 2 h before addition of IFN-β and then cultured for 48h (followed by a 6 to 8 h resting phase in cytokine-free media). Their ability to respond to IL-10 (40 ng/ml) or IL-6 (40 ng/ml) was then measured by means of P-STAT3 levels assessed via Phospho-Flow. (**B**) Schematic representation of the experimental approach adopted to assess the direct or indirect impact of IFN-β. (**C**) Purified T cells from congenic C57BL/6 (CD45.1) and IFN-AR1^−/−^ (CD45.2) mice were mixed at a 1:1 ratio.This mix was then cultured for 48h with or without IFN-β (5ng/ml), and rested in cytokine-free media for additional 6 h. The response to IL-10 of each sub-population was then assessed via Phospho-Flow. Data of n=3 individual experiments are shown and expressed as % of Ratio MFI±SEM, *p<0.05, **p≤0.01, paired Student’s t test.

Our data indicate that TI-IFN causes inhibition of IL-10 signaling through a process that requires 24/48 hours of exposure. This suggests that multiple intracellular molecular modifications are needed to achieve this phenotype and the process could require the synthesis and activity of additional extracellular mediators. To test this hypothesis, we employed a co-culture system with T cells deficient for the receptor of TI-IFN (IFN-AR1^−/−^; unable to respond to IFN-α or -β).These cells have a response to IL-10 comparable to that displayed by wild type cells (Supplementary Fig. 4). We then exposed B6 wild type congenic T cells (CD45.1^+^ from B6/SJL mice) mixed at 1:1 ratio with B6-IFN-AR1^−/−^ T cells (expressing the CD45.2 isoform) to IFN-β. If the TI-IFN-induced inhibition of IL-10 signaling requires the synthesis and secretion of an intermediate factor, this molecule would also affect the response of IFN-AR1^−/−^ T cells in our co-culture system (Fig. 5B). Phospho-flow analysis indicated that while wild type Tmem and Treg cells showed a reduction of IL-10 signaling after IFN-β exposure, the co-cultured IFN-AR1^−/−^ cells remained completely unaffected (Fig. 5C). These results demonstrate that IFN-β acts directly on T cells to condition IL-10 signaling.

### 2.6 Alteration of surface IL-10 receptor expression or induction of SOCS molecules are not responsible for inhibition of IL10 signaling

Downregulation of IL-10R surface expression would be a plausible mechanism to account for the inhibition of IL-10 signaling following exposure to TI-IFN. However, flow cytometric analysis of IL-10R surface expression did not support this hypothesis. IFN-β exposed T cells (both Tmem and Treg), expressed levels of the receptor comparable to that of non-exposed cells (Fig. 6A). In line with this, the comparison of IL-10R expression between NOD T cells isolated from the PLN, MLN, ALN, and spleen showed no differences in the mean fluorescence intensity (Fig. 6B). This result suggested the involvement of an IL-10R specific regulator acting between the receptor and STAT3 (as STAT3 remains available for the IL-6 receptor to be phosphorylated). Suppressor of cytokine signaling proteins (SOCSs) act as cytokine-inducible negative regulators of cytokine signaling. We tested if the transcriptional levels of SOCS1 (the only regulator reported in the literature to be associated with inhibition of IL-10 signaling in a lymphoma cell line) [29] and SOCS3 (as control) were increased by IFN-β treatment. Despite a reduction in the SOCS1 and SOCS3 RNA levels in cultured T cells compared to fresh cells, the transcriptions levels of SOCS1 and SOCS3 were not increased by IFN-β treatment (Fig. 6C).These results were confirmed in Tmem and Treg subpopulations via Prime FlowRNA (Affymetrix) - technology that allows detection via flow cytometry of RNA and protein expression simultaneously at single cell level - clearly showing that the levels of SOCS1 and SOCS3 RNA were not up-regulated in Tmem and Treg cells exposed to IFN-β (Fig. 6D).

**Figure 6.**
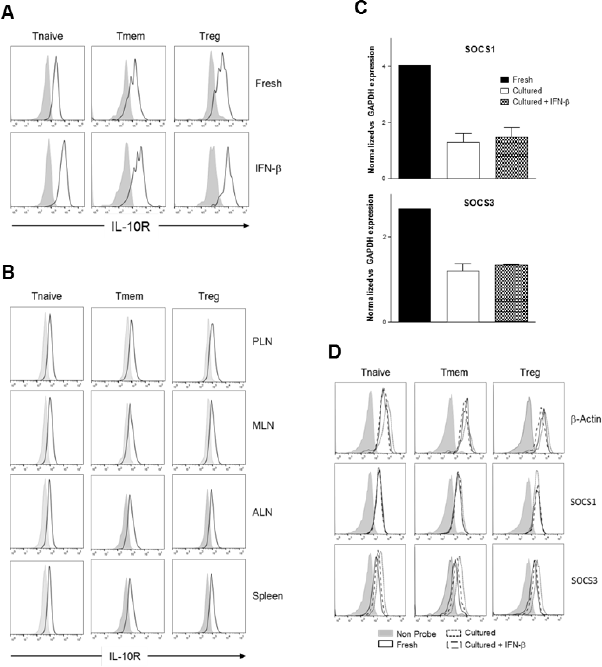
Alteration of surface IL-10R expression or induction of SOCS molecules are not responsible for the inhibition of IL-10 signaling induced by IFN-β. (**A&B**) T cells from C57BL/6 mice (freshly-isolated or post in vitro exposure to IFN-β; A) or total cells from PLN, MLN, axillary LN (ALN), and spleen of 4 week old NOD mice (B) were stained for their expression of IL-10R in the indicated CD4 T cell subpopulation by flow cytometry. (**C&D**) Purified CD4 Tmem (obtained via the “parking method) and Treg (C), or bulk T cells from C57BL/6 mice (D) were left untreated or cultured for 48h with or without IFN-β (5ng/ml) and the mRNA levels of SOCS1 and SOCS3 assessed by qPCR (C) and by Prime FlowRNA (D). In C, graph bars show qPCR results, calculated using ΔΔCt method and normalized with GAPDH expression. Data of n=3 individual experiments are shown and expressed as a Fold change ±SEM.In D, histograms show mRNA levels of SOCS1 and SOCS3 in CD4 T cell subpopulations in the indicated conditions; levels of β-actin are also shown as a control. Representative results of n=2 experiments are shown.

### 2.7 The localized defective response to IL-10 in Tmem and Treg cells of NOD mice is maintained with age

Knowing if the inhibition of IL-10 signaling in T cells of PLN and MLN of NOD mice is sustained or altered over time is important for defining possible windows of therapeutic intervention. The accumulation of TI-IFN in PLN [13] appears maintained over time, though to a progressively lower level. As the inhibition of IL-10 signaling correlates with TI-IFN levels, we executed a longitudinal analysis of the response to IL-10 and IL-6 by Treg and Tmem cells of female NOD mice of different ages (weeks: 2, 3, 4, 6, 12 and 18). An identical analysis was performed in age-matched female wt B6 mice. Throughout the observation period, the ratio of IL-10 induced P-STAT3 in T cells between PLN/spleen and MLN/spleen (as shown in Fig. 1) was significantly lower than that exhibited by B6 mice (Fig. 7A). Interestingly, in 2 week old B6 mice the response to IL-10 of Tmem and Treg cells in PLN and MLN was slightly lower than that observed in the spleen (<100%), but it quickly recovered and stabilized (Fig. 7A).

**Figure 7.**
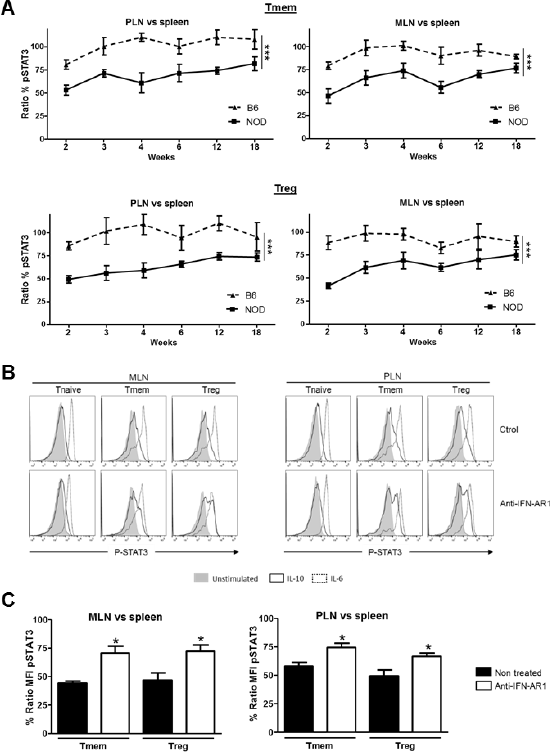
The localized inhibition of IL-10 signaling in T cells is sustained throughout the life of NOD mice and depends on IFN-β signaling. (**A**) P-STAT3 induction in response to IL-10 in T cell subsets of PLN, MLN, and spleen collected from NOD and C57BL/6 mice at the indicated age. Graph indicate the kinetic of P-STAT3 MFI ratio between the spleen and PLN or MLN (as indicated) Tmem and Treg after IL-10 stimulation. Final value was calculated by comparing the coefficient index of P-STAT3 between the two different organs, considering the levels in spleen as 100% of expression. Data of n=4 mice per strain and per age point are shown and expressed as % of Ratio MFI±SEM. Differences between B6 and NOD were calculated using a Two-Way ANOVA ( ***p<0.001). (**B&C**). Transient blockade of TI-IFN signaling partially restores IL-10 signaling in T cells. NOD mice were treated with control or a blocking anti-IFNA-R mAb (0.5 mg on day 14 and 21 of age) and, on day 25, PLN, MLN and spleen were extracted. Cells were left untreated or stimulated with IL-10 (40 ng/ml) or IL-6 (40 ng/ml) and the levels of P-STAT3 in CD4 T cell subpopulations measured via Phospho-flow. Representative histograms are presented in (B). Cumulative results are shown in (C), where graph bars represent the normalized P-STAT3 MFI ratio in Tmem or Treg after IL-10 stimulation between the spleen and PLN or MLN in control vs treated NOD mice. Ratio MFI was calculated by comparing the coefficient index of P-STAT3 between the two different organs, considering the levels in spleen as 100% of expression. Data of n=4 mice per treatment group are shown and expressed as % of Ratio MFI±SEM, *p<0.05, paired Student‘s t test.

### 2.8 The early blockade of TI-IFN signaling partially restores the response to IL-10 in PLN and MLN Tmem and Treg cells in NOD mice

The reported delay and lower incidence of diabetes development in NOD mice treated at 2-3 weeks of age with an anti-IFN-AR1 mAb [13], together with the reversibility of IL-10 signaling inhibition we observed in vitro in T cells after removing IFN-β, are encouraging indications that the unbalanced immune regulation of NOD mice can be prevented or restored. To test if the therapeutic blockade of TI-IFN and alteration of IL-10 signaling were correlated *in vivo* in NOD mice, anti-IFN-AR1 mAb was administered on days 14 and 21 of age in female NOD mice and, on day 25, we measured the induction of P-STAT3 in PLN and MLN T cell subsets in response to ex-vivo IL-10 treatment. Both Tmem and Treg populations in PLN and MLN of anti-IFN-AR-treated animals showed a significant recovery in P-STAT3 induction in response to IL-10 (Fig. 7B&C). These results suggest that the therapeutic effect of early administration of anti-IFN-AR1 mAb in NOD mice may be due to the restoration of IL-10 signaling.

## 3. Discussion

In addition to genetic susceptibility, type 1 diabetes development is clearly linked to still poorly defined environmental factors contributing to the initiation and/or progression of the disease. There is a growing number of reports that associate high levels of TI-IFN with the onset and progression of this pathology [8–10, 12–14]. However, mechanistic insights on how TI-IFN would favor type 1 diabetes development are lacking, contributing to a degree of confusion in understanding this connection. In particular, how the accumulation of TI-IFN could result in a localized effect that enables the activity of diabetogenic T cells remains an enigma. The finding that dendritic cells of NOD mice produce higher levels of TI-IFN in response to TLR stimulation than cells of B6 mice [30] provides an important clue to the causes of localized and chronic TIIFN accumulation in PLN reported previously [13]. Based on our results, we propose that this chronic accumulation of TI-IFN in NOD mice, which we report for the first time also extends to the MLN, re-programs the response of CD4 Treg and Tmem to IL-10 and potentially diminishing the homeostatic regulation of diabetogenic cells. This phenomenon likely contributes to the progressive attack to the islets and development of the disease. This data is supported by the recovery of IL-10 signaling we observed after blockade of TI-IFN-IFNR1 interactions, a treatment that results in a reduction of the disease incidence and progression in NOD mice [13].

Our results provide some important clues on the molecular mechanism behind the inhibition of IL-10 signaling. The normal level of phosphorylation of STAT3 we measured in response to IL-6 in T cells pre-exposed to IFN-β suggested that the impairment observed in IL-10 signaling is not caused by a generalized reduction in the cytoplasmic availability of STAT3. The lack of upregulation of mRNA levels of SOCS1 and SOCS3 and the unaltered (if not slightly increased) expression of IL-10R in T cells we observed post-IFN-β exposure both in vitro and in vivo do not support regulation of P-STAT3 induction at this level. Our results suggest then that an IL-10 signaling negative regulator acting between the receptor and STAT3 phosphorylation is either being upregulated or activated. For example, direct or indirect alterations in the binding and phosphorylation of Jak1 and Tyk2 would regulate STAT3 activation [31]; the involvement of molecules that interact with STAT3, including other members of the STAT family, should also be explored as they could interfere with its binding to IL-10R and its phosphorylation [28].Interestingly, human monocytes and macrophages primed with IFNα/β showed an increase in IL-10R1 and an increase in IL-10 signaling [32]. However, these experiments were performed with an acute exposure (5 hours) to TI-IFN. A prolonged exposure of human macrophages to TIIFN promoted a switch in the signaling of IL-10 from activation of STAT3 to STAT1 [33]; in our settings however STAT1 phosphorylation was not appreciable in any condition tested (not shown). Overall, these observations suggest that IFN-β can have different consequences depending on the timing and context of exposure as well as on the target cell population, probably contributing to the (sometimes discordant) range of outcomes attributed to TI-IFN [34].

The protective effect of Tofacitinib and Ruxolitinib we observed in our experiments indicates that signaling through the Jak/STAT pathway is involved in this specific effect of TI-IFN – though we cannot exclude the participation of other signaling routes [35]. A deeper understanding of the molecular mediators of this phenomenon would be of crucial importance as some of these molecules could be targeted to control the development of diabetes. In fact, Jak1/Jak2 inhibition in vivo is effective at preventing, and reverting, established insulitis in NOD mice [36, 37]. Improving the efficacy and safety of this type of intervention would be a major advancement for type 1 diabetes patients.

We observed that the impact of TI-IFN is not specific to the NOD genetic background, suggesting that this novel mechanism of alteration of immunoregulation could also contribute to the development of other disorders with a TI-IFN signature.[40] This is a significant finding as the reduction of IL-10 signaling in Treg and Tmem populations has very deleterious effects on regulating autoreactivity [22, 24, 25]. In animal models of diabetes [41, 42] as well as diabetic patients [43], regulatory T cells exhibit reduced regulatory efficacy while effector T cells are resistant to regulation [42]. Altogether these results point toward a loss of regulation of autoreactive T lymphocytes as a key process in diabetes development. A selective targeting of this phenomenon, by either correcting the aberrant production of TI-IFN or by preventing its modulation of IL-10 signaling, could significantly improve the efficacy of some approaches currently being explored for the treatment of type 1 diabetes [44]. Targeting IL-10 signaling has already been considered in the treatment of this disease [16, 17]. However, inconsistent effects have been reported [45–48]. Based on our results, we suggest that, rather than augmenting the concentration of IL-10, a targeted restoration of the IL-10 impact on diabetogenic cells (via a timed and localized intervention) would achieve more successful therapeutic outcomes. To this end, understanding how IL-10 modulates T cell function is necessary to identify the best strategy to recover an appropriate level of regulation but, to date, very little is known. We report here for the first time several genes (*Sphk1, LIGHT, Tarm1 and 2B4*) that IL-10 induces in T cells. We used their expression as readout of IL-10 function, demonstrating the impact of pre-exposure to TI-IFN. Future studies will be needed to understand their (and others‘) involvement in the modulation of T cell functions. Moreover, their differential expression profile between Tmem and Treg populations, suggests a distinct role in each population: a property that could also reveal strategies to selectively impact these two subsets.

Our study shows that Tmem and Treg with impaired IL-10 signaling are present not only in PLN but also in the regionally close MLN. Paralleling the results from Rahman and colleagues [30], this phenomenon is specific for NOD animals, as in B6 mice, the response to IL-10 was not affected. This strain-specific effect in lymph nodes draining the gut [38], suggests a link with the recently discovered role of the microbiota in autoimmune diabetes development [3, 4]: it could indicate an aberrant heightened chronic response to specific bacteria (or viruses) derivatives that is not properly regulated. The initial impairment in IL-10 signaling we observed in two-week-old B6 pups (similar to, but not to the same extent as, that of NOD mice, Figure 7), could indicate an initial adaptation phase of the newly generated pool of T cells (exposed to TI-IFN in the thymus) [39] to variations in the intestinal flora during the breast-feeding phase. This would suggest that the genetic predisposition of NOD mice encompass a defect in establishing the proper balance in the response to microbiota derivatives in the gut-draining lymph nodes that ultimately affects the regulation of diabetogenic T cells by IL-10.

In summary, our study unveils the existence of a new molecular mechanism through which TI-IFN can alter T cell regulation and improves our understanding of IL-10 mediated control of Treg and Tmem cells. A deeper understanding of this phenomenon will very likely reveal novel points of intervention to restore the necessary immune regulatory network to potentiate the efficacy of immunotherapies for type 1 diabetes and possibly other autoimmune diseases.

## 4. Methods

### 4.1 Mice

NOD, wt C57BL/6 (B6), IFN-AR1^−/−^ and Rag^−/−^ mice were purchased from Jackson Laboratories, and bred at the Johns Hopkins School of Medicine facility. All animal experiments were conducted in accordance with the National Institutes of Health guide for use and care of laboratory animals, and under a protocol approved by the JHU Animal Care and Use Committee.

### 4.2 Media, reagents and antibodies

RPMI-1640 and IMDM media were supplemented with 10% v/v heat-inactivated FCS (Atlanta Biologicals, Flowery Branch, GA), 0.1 mM non-essential amino acids, 2 mM L-glutamine, sodium pyruvate, 100 IU/ml penicillin, 100 µg/ml streptomycin, and 50 µM 2-ME (Gibco).Recombinant IFN-β was purchased from PBL Assay Science. Blocking anti-IFN-AR1 mAb was from Leinco Technologies (St Louis, MO). Recombinant IL-10, and IL-6 were from PeproTech (Rocky Hill, NJ). Jak inhibitors Tofacitinib and Ruxolitinib were purchased from LC Laboratories (Woburn, MA).

### 4.3 T cell (subsets) isolation

Spleen and lymph nodes were harvested and total/CD4 T cells were isolated via magnetic-bead negative selection. Briefly, cells were incubated with: anti-mouse Ter119 (TER-119), Gr1 (RB6-85C), CD11b (M1/70), B220 (RA3-6B2), CD16/32 (2.462), I-A/I-E (M5/114.15.2), (also anti-CD8 (53-6.7) for CD4 T cell purification) (all from BD Biosciences) followed by incubation with magnetic beads conjugated with anti-rat IgG (ThermoFisher) at a 1:1 (cell:bead) ratio. The resulting total/CD4 T cells were >90% pure. Where indicated, Treg (CD4^+^CD25^+^) were isolated from CD4 T cells following the protocol described in the EasySep PE-selection kit (STEMCELL technologies).

### 4.4 CD4 memory T cell generation

In some experiments, memory T cells were generated via a modification of the published “parking method” [27]. Briefly, 20×10^7^ T cells from B6 mice were activated with anti-CD3 (0.5 µg/ml; BD Pharmingen) in the presence of syngeneic LPS-matured bone marrow-derived DCs (1:20 ratio DC:T cell) as previously described [49]. Three days later, activated T cells together with 10^7^ T cell-depleted splenocytes (obtained via removal of CD3+ cells from single cell suspensions using the protocol described in 2.3) were infused intravenously into Rag^−/−^ mice. Four weeks later, memory T cells were isolated and used for the indicated experiments.

### 4.5 Cell stimulation and preparation for Phospho-flow analysis

For assessment of proteins phosphorylation via flow cytometry (Phospho-flow) a modification of the protocol published by the Nolan group [50] was utilized. Briefly, 10 to 15×10^6^ purified T cells were cultured in IMDM complete media with/without IFN-β (1-25 ng/ml) for indicated periods and then rested in cytokine-free media for six additional hours. 2×10^6^ fresh/cultured cells were stimulated for 20 minutes with IL-10 (40 ng/ml) or IL-6 (40 ng/ml), or 30 minutes with IFN-β (5 ng/ml). Then, cells were fixed for 50 minutes by adding 2.4 ml of a solution containing 4% paraformaldehyde and 1.4% methanol in PBS. After fixation, 600 µl of 1X wash buffer (contained in the Transcription Factor Phospho Buffer Set kit, BD Biosciences) were added to the previous mixture, mixed, and spun down. Finally, cells were suspended with 500 µl Perm Buffer III (BD Biosciences) while vortexing, and stored at −20^o^C until use.

### 4.6 Flow cytometry

In Phospho-flow experiments, Perm Buffer-III was removed and cells stained with fluorchrome-labeled antibodies against CD4 (RM4-5), CD44 (IM7), Foxp3 (FJK-1) (Thermo Fisher eBioscience) and Stat3 (pY705)(4/P_STAT3; BD Phospho-flow). For IL-10R staining, the Human IL-10 biotinylated Fluorokine kit (R&D Systems) was used. Detection of SOCS1 and SOCS3 mRNA levels via flow cytometry was performed employing the PrimeFlow RNA assay kit (Thermo Fisher) following manufacturer guidelines. Data were acquired using a LSR-II flow cytometer (BD Biosystems) and analyzed with FlowJo X version software (FLOWJO, LLC, Ashland OR).

### 4.7 Quantitative Real-Time PCR

CD4 T cells were lysed in TRIzol reagent (Thermo), and RNA was extracted using chloroform (Fisher Scientific) and the RNeasy MiniElute Cleanup Kit (Qiagen). The mRNA was reverse transcribed using the SuperScript IV First Strand Synthesis System and protocol (Thermo Scientific). Real time RT-PCR was performed on a QuantiStudio 12K Flex Real Time PCR system (Thermo Scientific) calibrated for SYBR Green detection. The primers and conditions employed are listed in Supplementary Table 1.

### 4.8 Statistical Analysis

Differences in flow cytometry quantification of P-STAT3-mean fluorescence intensity (MFI) were analyzed using two-tailed paired Student *t* test. To minimize the impact of fluctuations in fluorescence among experiments, the coefficient index (MFI P-STAT3 in stimulated cells/MFI P-STAT3 in unstimulated cells) was calculated and averaged for each experiment and then used for statistical analysis. P-STAT3 expression comparisons between T cells of NOD and B6 mice were analyzed using Two-way ANOVA. Two-tailed unpaired Student *t* test was applied to test gene expression differences from PCR experiments. All analyses were performed with Prism Software version 5.0 (GraphPad, La Jolla, CA).

## Aknowledgements

The authors thank Sonia Santiago (laboratory manager), Xiaoling Zhang (flow cytometry specialist), both at Johns Hopkins University, for excellent technical assistance; Dawn Hull and Kamal Abdi (animal facility supervisors; Johns Hopkins University) for animal husbandry and care. We would also like to thank Dr. Ranjeny Thomas (University of Queensland, Australia), Dr. John Alcorn (University of Pittsburgh), Drs. Alan Scott, Thomas Donner, Edward Harhaj, Erika Darrah, Tory Johnson (Johns Hopkins University), Dr. Francesca Granucci (University of Milano-Bicocca, Italy), and Dr. Ivan Zanoni (Harvard University) for invaluable feedback on experimental design, troubleshooting, and manuscript preparation.

## Author contributions

M.I and G.R. developed the project, researched data, designed experiments, and wrote the manuscript. A.A., M.C, and B.L. contributed to experimental design and researched data. C.T. and V.I. researched data. W.P.A.L and G.B. contributed to data interpretation, troubleshooting, and provided essential manuscript feedback. G.R is the guarantor of this work and, as such, had full access to all the data in the study and take responsibility for the integrity of the data and the accuracy of the data analysis.

## Supplementary Figures

**Figure Supplementary 1.**
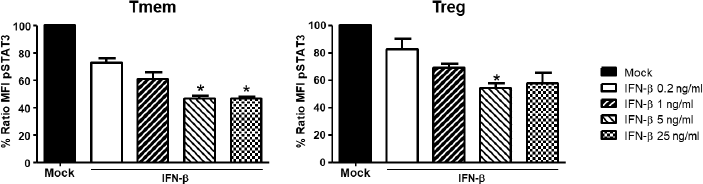
Dose-effect of IFN-β inhibiting IL-10 signaling in T cells. Purified T cells from C57BL/6 mice were cultured for 48h in RPMI complete media in the absence (mock culture) or presence of different concentrations of IFN-β (0.2-25ng/ml), and rested in cytokine-free media for six more hours. Cells then were either left untreated or stimulated with IL-10 (40 ng/ml) for 20‘. After a fixation step, the levels of P-STAT3 in CD4 T cell subpopulations (Tmem: CD4^+^CD44^hi^Foxp3^-^, Treg: CD4^+^Foxp3^+^) were measured by phosphor-flow. The graph bars compare the percentage of P-STAT3 MFI ratio in Tmem and Treg cells between the groups exposed to IFN-β and mock condition (considered as 100% of response) after IL-10 stimulation. Ratio MFI was calculated comparing the coeficient index of P-STAT3 after stimulation between each of the IFN-β exposed groups and mock. Data of n=3 individual experiments are shown and expressed as % of Ratio MFI±SEM, *p<0.05, paired Student‘s t test.

**Figure Supplementary 2.**
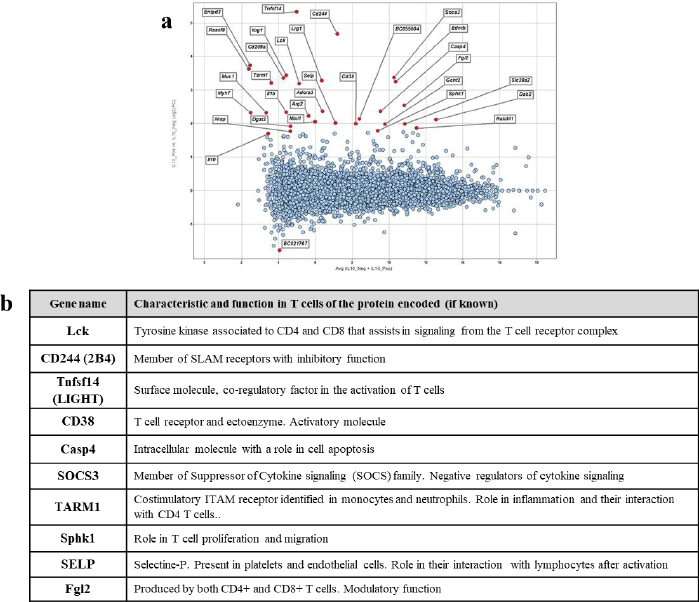
Genes tested for expression/modulation after IL-10 stimulation in T cells. (a) Re-analysis of the RNAseq project evaluating mouse macrophage expression in response to IL-10 stimulation, GSE49449, deposited in the NCBI Gene Expression Omnibus (GEO) database. A two-tailed one-way ANOVA was performed to determine IL-10-treated versus untreated cells’ differential gene expression. This differential expression was then evaluated for the standard deviation, σ or SD, of each gene‘s log2 fold change from the mean, of zero or unchanged. The 29 genes highly upregulated (>6SD) are indicated. (b) The table shows the 10 genes among the 29 indicated in (a), that we found also described in T lymphocytes. Characteristics and function (if known) of the protein encoded are indicated.

**Figure Supplementary 3.**
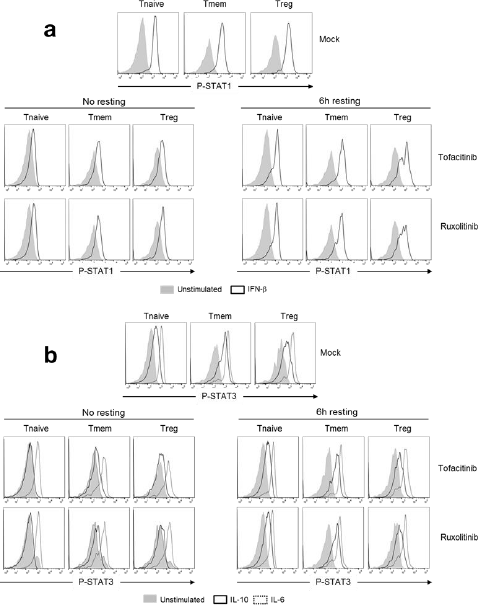
Effect of Tofacitinib and Ruxolitinib in cytokine signaling. (a&b) Purified T cells from C57BL/6 mice were cultured for 48h in RPMI complete media with or without Tofacitinib (25μM) or Ruxolitinib (5μM), and then directly stimulated or rested in cytokine-free media for additional 6 h. Cells in every of the mentioned conditions were either left untreated or stimulated with IFN-β for 30 min (A) or with IL-10 (40 ng/ml) or IL-6 (40 ng/ml) for 20 min (B). Representative histograms show the levels of P-STAT-1 (A) or P-STAT3 (B) after stimulation in CD4 T cell subpopulations (Tnaive: CD4^+^CD44^low^Foxp3^-^, Tmem: CD4^+^CD44^hi^Foxp3^-^ and Treg: CD4^+^Foxp3^+^) after stimulation in the different conditions. Results are representative of n=3 independent experiments.

**Figure Supplementary 4.**
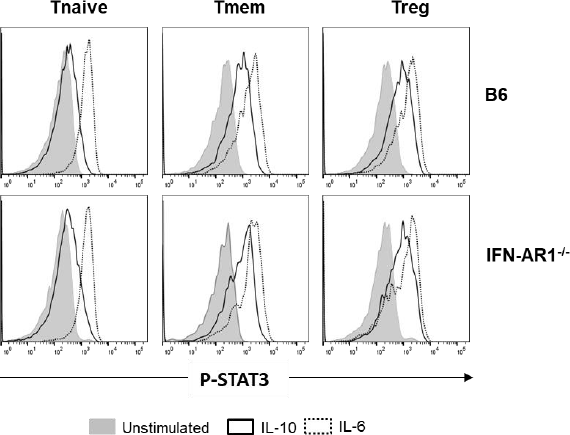
Response of IFN-AR1^−/−^ T cells to IL-10 stimulation. Freshly purified T cells from C57BL/6 and IFN-AR1^−/−^ mice were either left untreated or stimulated with IL10 (40 ng/ml) or IL-6 (40 ng/ml) for 20‘. The levels of P-STAT3 after stimulation in CD4 T cell subpopulations were measured by phosphor-flow. Representative histograms show P-STAT3 levels in Tnaive: CD4^+^CD44^low^Foxp3^-^, Tmem: CD4^+^CD44^hi^Foxp3^-^ and Treg: CD4^+^Foxp3^+^ cells after stimulation in both cell types. Results are representative of n=3 independent experiments.

## Supplementary Tables

**Table Supplementary 1.**
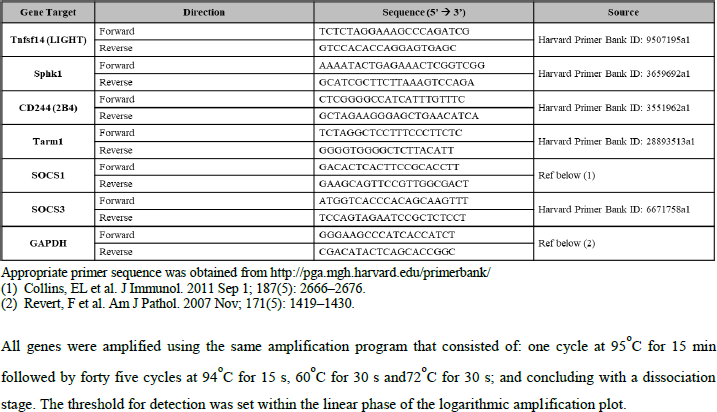
Sequence of primers used in RT-PCT studies.

